# Soil-derived *Bacillus pumilus* strains demonstrate antagonistic activity against *Magnaporthe oryzae* and multiple plant growth-promoting traits

**DOI:** 10.64898/2026.06.28.735134

**Authors:** Lainey E. Kemmerer, Timothy R. Johnson, Garrett L. Ellward, Rachel E. Kalicharan, Nalleli Payne, Daniel M. Czyz, Jessie Fernandez

## Abstract

Biological control strategies are increasingly being explored as sustainable alternatives for managing rice blast disease caused by *Magnaporthe oryzae*. In this study, we characterized three *Bacillus pumilus* isolates (DC01, DC09, and DC13) and evaluated their antifungal and plant-beneficial properties against *M. oryzae*. Whole genome sequencing revealed multiple biosynthetic gene clusters associated with the production of antimicrobial metabolites. All three isolates inhibited fungal growth in dual-culture assays, whereas heat-stable diffusible antifungal activity was primarily associated with the cell-free supernatants of DC09 and DC13. Exposure to bacterial supernatants disrupted fungal development, inducing abnormal hyphal morphology characterized by bulbous swelling, altered polarity, and increased branching in *M. oryzae*. Volatile organic compound assays further revealed that the DC isolates suppress fungal growth in the absence of physical contact. The isolates additionally inhibited the growth of other phytopathogenic fungi and selected human bacterial pathogens. All strains exhibited plant growth-promoting traits, including indole-3-acetic acid production and osmotic stress tolerance, whereas DC09 also displayed phosphate-solubilizing activity. Importantly, root inoculation with the DC isolates significantly reduced rice blast disease severity and induced expression of defense-associated genes involved in jasmonic acid/ethylene signaling and immune priming. Collectively, these findings identify the DC isolates, particularly DC09 and DC13, as promising multi-mechanistic biological control agents for sustainable rice blast management.

## Introduction

Rice blast, caused by the filamentous fungus *Magnaporthe oryzae*, is one of the most significant threats to global rice production, causing annual yield losses of 10-30% [1, 2]. The pathogen infects rice through specialized appressoria that generate high turgor pressure to penetrate host tissues, followed by colonization and prolific sporulation that facilitates disease spread [1, 3, 4]. This highly efficient infection cycle, combined with adaptability and reproductive capacity, underlies its epidemic potential and persistent threat to global food security [5, 6].

Management of rice blast relies primarily on chemical fungicides and resistant cultivars. However, concerns regarding environmental impact, resistance development, and the limited durability of resistance genes reduce the long-term effectiveness of these approaches [7–9]. These limitations highlight the need for complementary and sustainable disease-management strategies.

Biological control using plant-associated bacteria has emerged as a promising strategy to enhance crop resilience and suppress disease. Many of these microorganisms, function as plant growth-promoting bacteria (PGPB), improving plant performance through mechanisms such as nitrogen fixation, phytohormone production, including indole-3-acetic acid (IAA), and phosphate solubilization [10]. They also suppress pathogens through the production of antimicrobial compounds, including lipopeptides and siderophores, and by triggering induced systemic resistance (ISR), which enhances host immunity [10, 11]. Through these combined activities, PGPB simultaneously promote plant vigor and reduce pathogen pressure.

Among PGPB, species within the genus *Bacillus* are particularly attractive for agricultural applications because of their environmental resilience and ability to produce diverse bioactive metabolites [12–14]. Several species, including *B. subtilis*, *B. amyloliquefaciens*, *B. licheniformis*, *B. megaterium*, *B. cereus*, *B. thuringiensis*, and *B. velezensis,* have been reported as effective PGPB and biocontrol agents across diverse cropping systems [15–19]. Their efficacy is largely attributed to the production of lipopeptides (surfactins, fengycins, iturins), polyketides, siderophores, and volatile organic compounds (VOCs), as well as their ability to induce ISR and persist under field-relevant conditions [11, 14].

Although less extensively characterized than other *Bacillus* species, *B. pumilus* is increasingly recognized for its potential as both a PGPB and a biocontrol agent [20]. Genomic analyses indicate that *B. pumilus* frequently harbors biosynthetic gene clusters (BGCs) encoding non-ribosomal peptide synthetases and polyketide synthases associated with secondary metabolite production [21, 22]. Several strains can colonize root and leaf tissues, form biofilms, and persist under fluctuating environmental conditions, highlighting traits that support both disease suppression and plant growth promotion. Importantly, individual *B. pumilus* strains often exhibit distinct subsets of PGP traits, including either IAA production, phosphate solubilization, nutrient fixation, stress tolerance, or induction of plant defense responses, rather than a single strain possessing all PGP properties [21, 22]. In addition, certain *Bacillus* strains trigger ISR through salicylic acid (SA)-, jasmonic acid (JA)-, and ethylene (ET)-regulated pathways, resulting in reduced lesion formation and enhanced defense gene expression in rice [20, 23–26]. Despite these promising qualities, the biocontrol and PGP potential of *B. pumilus* remains less explored than that of other *Bacillus* species.

In this study, we identified three *B. pumilus* strains recovered from topsoil samples in Gainesville, Florida, and performed an integrated assessment of their anti-fungal and plant-beneficial properties. We evaluated their antagonistic activity against *M. oryzae*, characterized PGP traits, and examined their ability to modulate host defense responses. Together, these findings identify the DC strains as promising multi-PGP-trait candidates for sustainable rice blast management.

## Results

### Genomic analyses identify the DC isolates as *B. pumilus* strains with diverse biosynthetic potential

To identify the bacterial isolates recovered from Gainesville soil samples, we sequenced the 16S rRNA gene using V3-V4 primers. All three isolates were identified as *Bacillus pumilus* and designated *B. pumilus* DC01, DC09, and DC13. Whole-genome sequencing further confirmed species-level identity. Draft genome assemblies ranged from approximately 3.7 Mb, with GC contents near 41%, values consistent with those of reference *B. pumilus* genomes. Assemblies for DC01 and DC09 exhibited high completeness (≥98%) and low contamination (<1%), whereas DC13 was recovered as a partial assembly. Nonetheless, GC content and phylogenetic placement supported its assignment to *B. pumilus*.

To assess biosynthetic potential, all genomes were analyzed with antiSMASH [27], using relaxed detection strictness and all features enabled (Table 1). Strains DC01 and DC09 exhibited highly similar biosynthetic profiles, harboring multiple BGCs associated with antimicrobial and metal-chelating metabolites, including predicted pathways for fengycin, schizokinen, bacilysin, lichenysin, and bacillibactin production. Both genomes also contained several unassigned terpene-, RiPP-like-, type III polyketide synthases (T3PKS)-, and β-lactone-associated regions. In addition, DC01 contained a sporulation killing factor-like Non-Ribosomal Peptide Synthetase (NRPS) cluster that was not identified in the other strains. In contrast, the partially assembled DC13 genome exhibited a more limited but distinct repertoire, including high- and medium-similarity clusters associated with bacillibactin and lichenysin, low-similarity clusters related to surfactin and zwittermicin A, and several unassigned β-lactone and RRE-containing loci. Because of the incomplete assembly, these clusters are likely to represent a minimum estimate of DC13’s biosynthetic capacity.

**Table 1.**
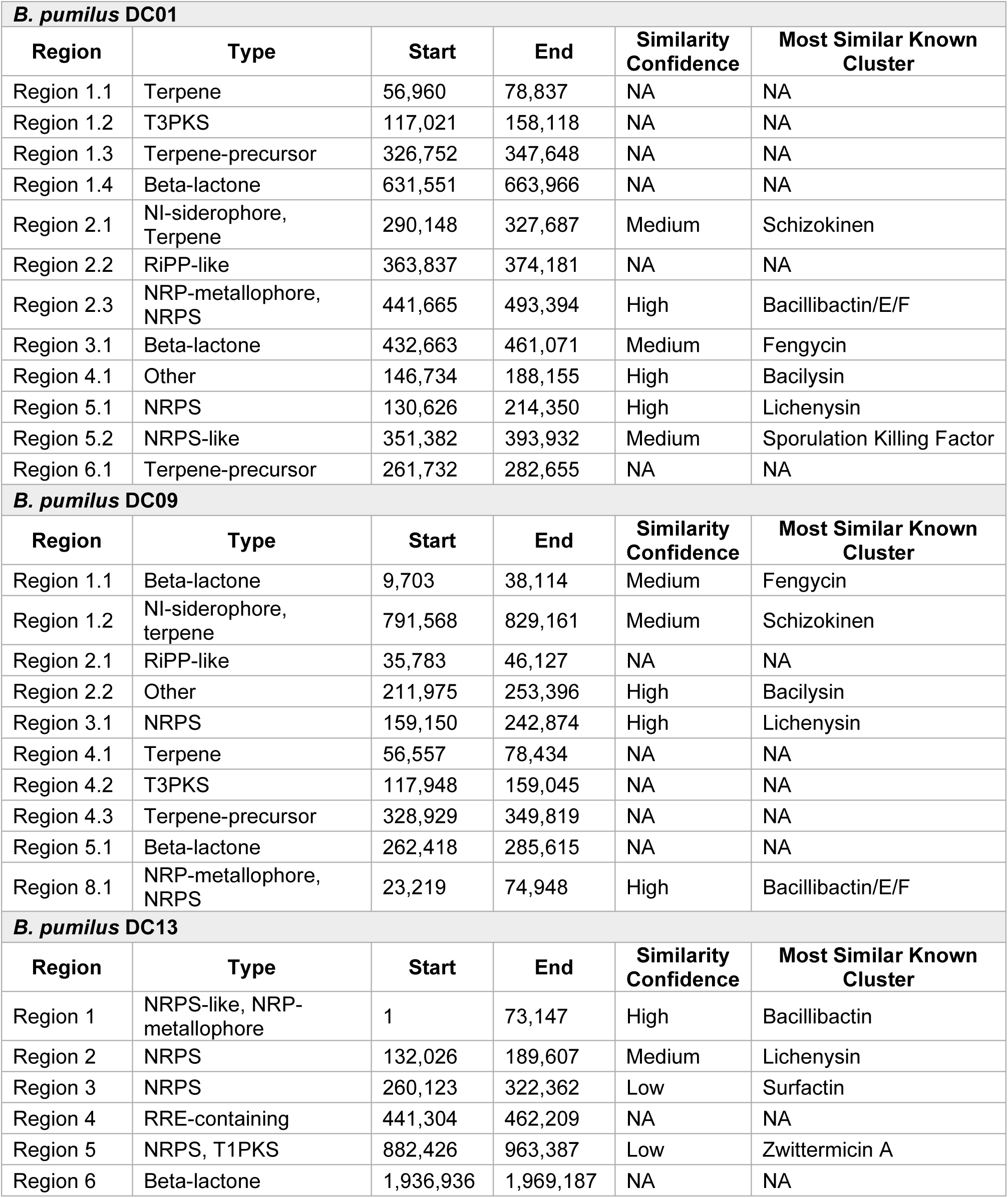
Biosynthetic gene clusters identified in AntiSMASH [27].

| ***B. pumilus* DC01** |  |  |  |  |  |
| --- | --- | --- | --- | --- | --- |
| **Region** | **Type** | **Start** | **End** | **Similarity Confidence** | **Most Similar Known Cluster** |
| Region 1.1 | Terpene | 56,960 | 78,837 | NA | NA |
| Region 1.2 | T3PKS | 117,021 | 158,118 | NA | NA |
| Region 1.3 | Terpene-precursor | 326,752 | 347,648 | NA | NA |
| Region 1.4 | Beta-lactone | 631,551 | 663,966 | NA | NA |
| Region 2.1 | NI-siderophore, Terpene | 290,148 | 327,687 | Medium | Schizokinen |
| Region 2.2 | RiPP-like | 363,837 | 374,181 | NA | NA |
| Region 2.3 | NRP-metallophore, NRPS | 441,665 | 493,394 | High | Bacillibactin/E/F |
| Region 3.1 | Beta-lactone | 432,663 | 461,071 | Medium | Fengycin |
| Region 4.1 | Other | 146,734 | 188,155 | High | Bacilysin |
| Region 5.1 | NRPS | 130,626 | 214,350 | High | Lichenysin |
| Region 5.2 | NRPS-like | 351,382 | 393,932 | Medium | Sporulation Killing Factor |
| Region 6.1 | Terpene-precursor | 261,732 | 282,655 | NA | NA |
| ***B. pumilus* DC09** |  |  |  |  |  |
| **Region** | **Type** | **Start** | **End** | **Similarity Confidence** | **Most Similar Known Cluster** |
| Region 1.1 | Beta-lactone | 9,703 | 38,114 | Medium | Fengycin |
| Region 1.2 | NI-siderophore, terpene | 791,568 | 829,161 | Medium | Schizokinen |
| Region 2.1 | RiPP-like | 35,783 | 46,127 | NA | NA |
| Region 2.2 | Other | 211,975 | 253,396 | High | Bacilysin |
| Region 3.1 | NRPS | 159,150 | 242,874 | High | Lichenysin |
| Region 4.1 | Terpene | 56,557 | 78,434 | NA | NA |
| Region 4.2 | T3PKS | 117,948 | 159,045 | NA | NA |
| Region 4.3 | Terpene-precursor | 328,929 | 349,819 | NA | NA |
| Region 5.1 | Beta-lactone | 262,418 | 285,615 | NA | NA |
| Region 8.1 | NRP-metallophore, NRPS | 23,219 | 74,948 | High | Bacillibactin/E/F |
| ***B. pumilus* DC13** |  |  |  |  |  |
| **Region** | **Type** | **Start** | **End** | **Similarity Confidence** | **Most Similar Known Cluster** |
| Region 1 | NRPS-like, NRP-metallophore | 1 | 73,147 | High | Bacillibactin |
| Region 2 | NRPS | 132,026 | 189,607 | Medium | Lichenysin |
| Region 3 | NRPS | 260,123 | 322,362 | Low | Surfactin |
| Region 4 | RRE-containing | 441,304 | 462,209 | NA | NA |
| Region 5 | NRPS, T1PKS | 882,426 | 963,387 | Low | Zwittermicin A |
| Region 6 | Beta-lactone | 1,936,936 | 1,969,187 | NA | NA |

Moreover, phylogenetic analysis using a codon-based tree constructed from 100 single-copy core genes and including 49 publicly available *B. pumilus* genomes placed the DC isolates firmly within the *B. pumilus* clade, forming a well-supported subgroup (100% bootstrap support) (Figure 1). Together, these genomic analyses confirm that the soil isolates are bona fide *B. pumilus* strains and reveal a diverse suite of BGCs associated with antimicrobial production and potential PGP functions.

**Figure 1.**
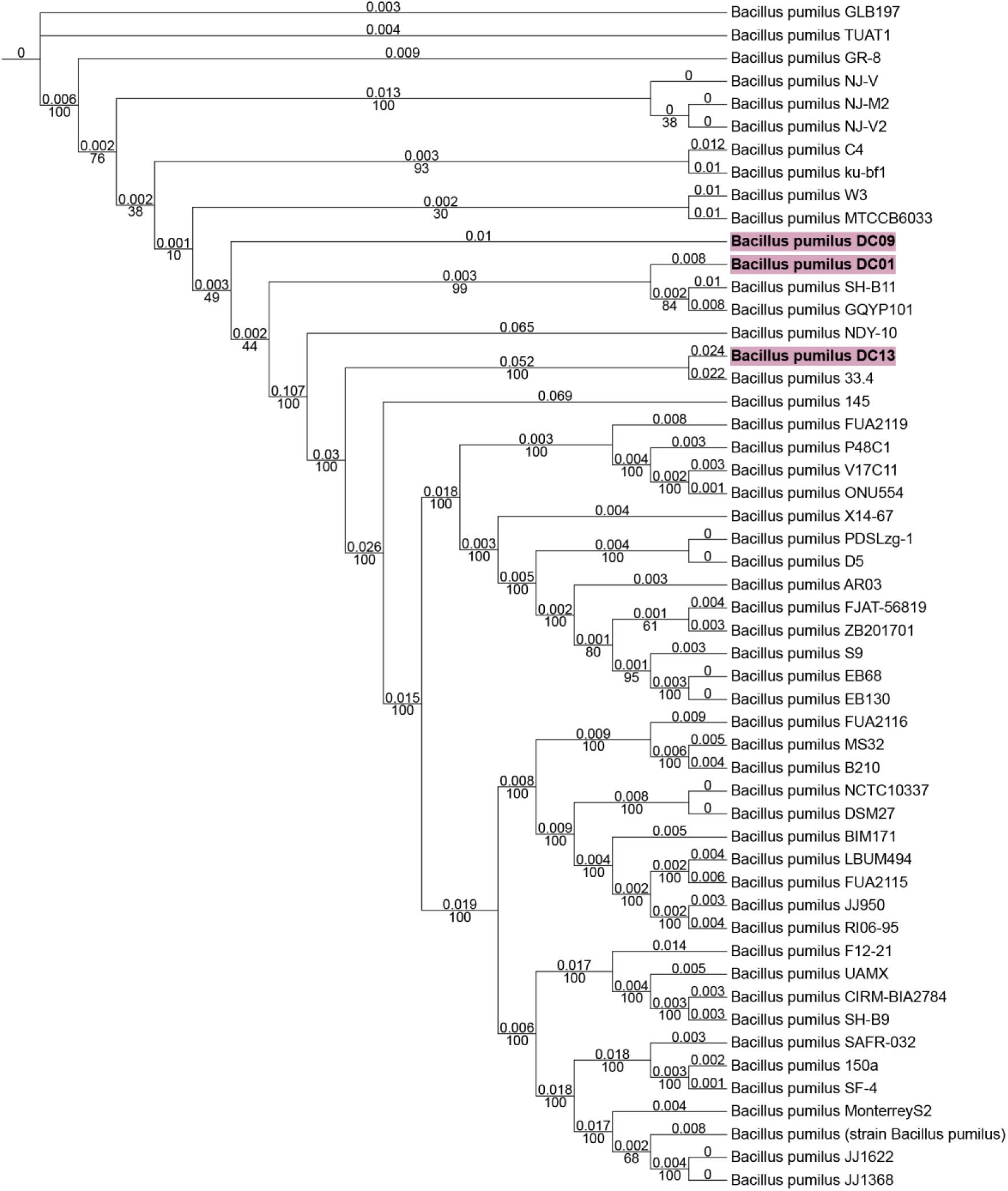
Phylogenetic analysis of DC isolates and publicly available *B. pumilus* genomes. A maximum-likelihood phylogenetic tree was generated from an alignment of 100 single-copy core genes from the DC isolates and 49 publicly available *B. pumilus* genomes using the BV-BRC phylogenetic pipeline. Core gene sequences were aligned with MAFFT, and the tree was inferred using RAxML. The tree was visualized and edited in iTOL where branch colors were added for clarity. Bootstrap values are shown at branch nodes. The scale bar represents the number of nucleotide substitutions per site. The DC isolates clustered within the *B. pumilus* clade, supporting their classification as *B. pumilus*.

### Soil-derived *B. pumilus* strains strongly inhibit *M. oryzae* growth *in vitro*

To assess whether the *B. pumilus* DC strains possess growth-inhibitory and antagonistic properties, we performed dual-culture assays on CM agar against *M. oryzae*. In this setup, each bacterial isolate (DC01, DC09, and DC13) was spotted opposite a central plug of the *M. oryzae* Guy11 strain, and fungal growth was monitored over a 10-day period (Figure 2A). At 10 days post-inoculation (dpi), all three isolates strongly suppressed fungal growth compared with uninoculated controls and the non-antagonistic control *E. coli* DH5α (Figure 2B). By 10 dpi, the overall colony diameter of *M. oryzae* was reduced by approximately 20-36% in the presence of DC01, DC09, or DC13 compared with untreated controls (Figure 2C). When fungal growth was quantified specifically along the axis facing the bacterial streak, inhibition was even more pronounced, reaching a 70-76% reduction in radial growth relative to mock-treatment plates (Figure 2D). At this stage, all three *B. pumilus* strains produced inhibition zones with a characteristic yellow-orange pigmentation at the bacterial-fungal interface. This pigmentation coincided with regions of visibly reduced hyphal density, suggesting local accumulation of antifungal metabolites within the inhibitory zone. These robust and reproducible antagonistic phenotypes supported the selection of DC strains for further characterization.

**Figure 2.**
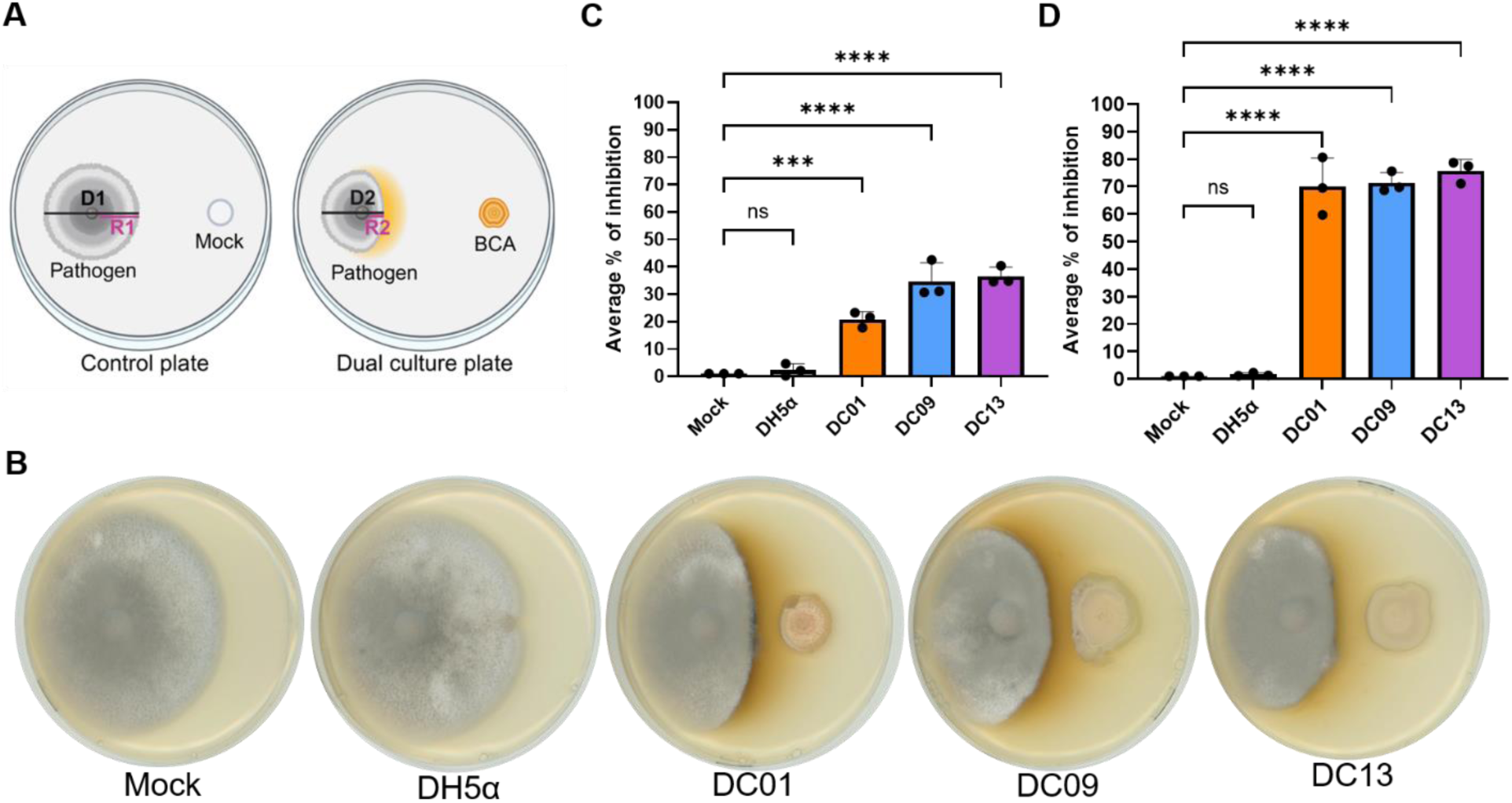
Antagonistic activity of *Bacillus* DC strains against *M. oryzae*. (**a**) Schematic diagram of the dual culture setup, where the control plate contains the pathogen with a water mock treatment, and the assay plate contains the pathogen with DC strains. D1 and D2 represent perpendicular measurements of the fungal colony diameter. R1 and R2 represent the mycelial radii measured from the point of inoculation toward the control mock and bacterial treatment sides of the plate, respectively. (**b**) Dual culture assay plates showing fungal growth inhibition by DC strains compared to controls (water and *E. coli* DH5α) at 10 days post-inoculation (dpi). (**c-d**) Percentage of fungal growth inhibition calculated from diameter (c) and radial (d) measurements, showing inhibition by DC strains relative to controls. Plates were incubated at 25 °C, and images and measurements were taken at 10 dpi. Bars represent mean ± standard deviation (SD) from three independent biological replicates (*n* = 3). Statistical analysis was performed using one-way ANOVA followed by Dunnett’s multiple comparison post-hoc test. ns (no significant), *p <* 0.001 (***), *p* < 0.0001 (****).

### Cell-free supernatants of *B. pumilus* strains inhibit *M. oryzae* growth

We next assessed whether cell-free supernatant (CFS) from the *B. pumilus* DC strains could inhibit *M. oryzae* growth *in vitro*, in the absence of living bacterial cells. Filtered CFS from DC01, DC09, and DC13, respectively, was incorporated into CM agar at a 1:6 ratio, and radial fungal growth was measured at 5 dpi. We observed that CFS from DC09 and DC13 significantly suppressed fungal colony expansion, whereas CFS from DC01 had little effect, comparable to the *E. coli* DH5α and mock controls (Figure 3). Quantitatively, DC13 CFS reduced radial growth by 62.6% ± 6.4%, and DC09 by 39.1% ± 6.4%. DC01 displayed only 2.5% ± 1.4% inhibition, which was not significantly different from the control treatments (Figure 3B). Similar inhibition patterns were also observed at 10 dpi (data not shown).

**Figure 3.**
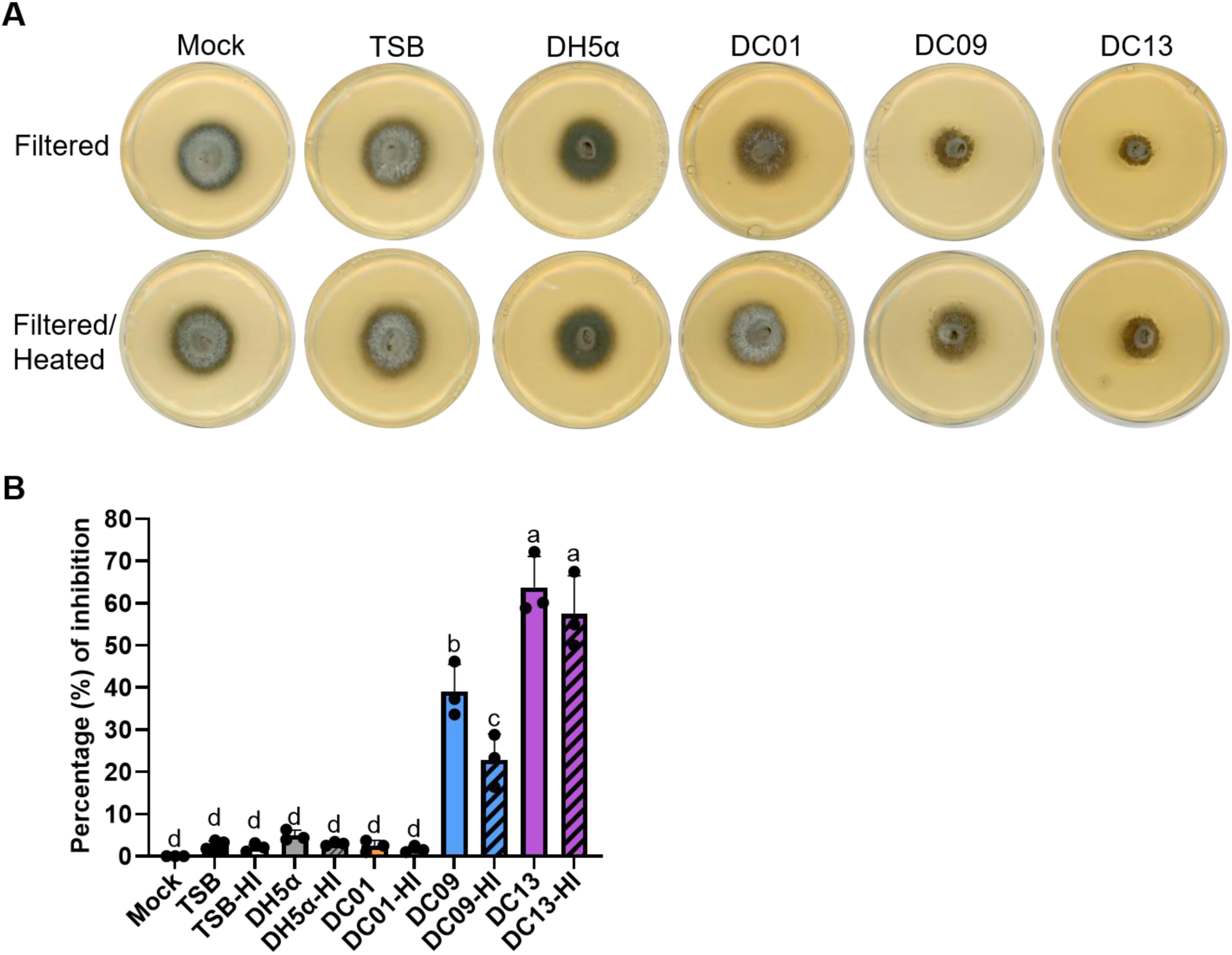
Growth inhibition of *M. oryzae* by cell-free supernatants of *B. pumilus* DC strains. (**a**) Representative CM agar plates showing fungal growth in the presence of cell-free supernatants (CFS) from DC01, DC09, and DC13, including heat-inactivated (HI) preparations. CFSs were incorporated into CM agar at a 1:6 ratio, and *M. oryzae* was point-inoculated at the center. Plates were incubated at 25 °C, and colony diameter was measured at 5 dpi. (**b**) Quantification of *M. oryzae* growth inhibition. Bars represent mean ± SD of three biological replicates. Statistical significance was assessed by one-way ANOVA followed by Tukey’s post hoc test for multiple comparisons among all treatments. Different letters indicate statistically significant differences among treatments (*p* < 0.05).

To test whether the antifungal factors are heat-sensitive, CFS samples were heat-inactivated (HI) prior to incorporation. After HI treatment, DC13-HI maintained a strong inhibitory effect (57.5% ± 9.0%), and DC09-HI retained moderate activity (22.8% ± 6.1%), both significantly greater (*p* < 0.05) than controls (Figure 3A, B). By contrast, DC01-HI showed negligible inhibition (1.6% ± 0.7%), similar to mock. These results indicate that DC09 and DC13 secrete stable antifungal metabolites that inhibit *M. oryzae* growth in a cell-free context.

To further examine how diffusible metabolites influence fungal development, *M. oryzae* was grown on thin CM agar layers supplemented with bacterial CFS and analyzed microscopically at 5 dpi. Hyphae exposed to DC isolate supernatants displayed sparse and irregular mycelial networks, with altered growth trajectories and reduced directional uniformity (Figure 4A). Among the treatments, DC13 produced the most pronounced effect, disrupting polarized hyphal extension and promoting extensive lateral branching. In contrast, hyphae grown on CM, CM supplemented with TSB, or CM containing *E. coli* DH5α supernatant exhibited dense and organized mycelial networks with continuous outward hyphal extension.

**Figure 4.**
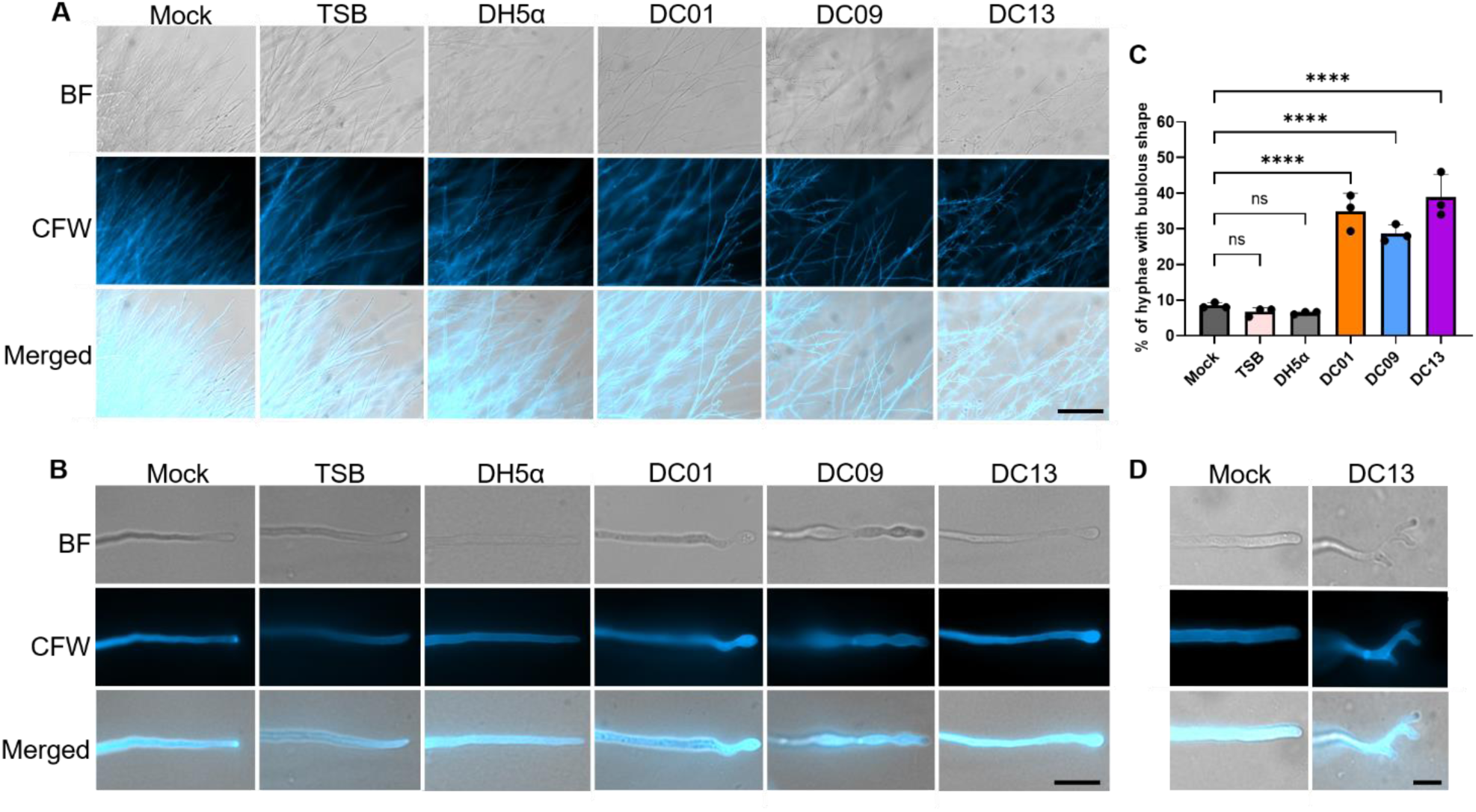
Effects of DC strains CFS on *M. oryzae* growth. (**a**) Brightfield, fluorescence, and merged images showing altered mycelial architecture and branching patterns following exposure to supernatants from DC01, DC09, and DC13, compared with mock, TSB, and DH5α controls. (**b**) Representative images of hyphal swelling and apical deformation induced by DC isolate supernatants. (**c**) Quantification of bulbous hyphae following treatment with the indicated supernatants. Data represent mean ± SD. Statistical significance was determined by one-way ANOVA followed by Dunnett’s multiple-comparison test relative to mock; \*\*\*\**p* < 0.0001; ns, not significant. (**d**) Representative images of hyperbranching and polarity defects observed following DC13 treatment. Scale bars: 100 μm (a), 10 μm (b), and 5 μm (d).

Visualization of individual hyphae further revealed that exposure to DC isolate supernatants induced hyphal swelling, resulting in a bulbous appearance at the apical regions (Figure 4B). DC01 and DC09 caused pronounced subapical swelling and localized tip abnormalities, whereas DC13 induced tip hyphal swelling and extensive branching defects, accompanied by apical curvature and subapical swelling (Figure 4B, D). Quantification of bulbous hyphae showed that DC01, DC09, and DC13 increased the proportion of abnormal hyphae to approximately 34.8% ± 5.0%, 28.6% ± 2.4%, and 38.9% ± 6.3%, respectively (Figure 4C). In contrast, mock, TSB, and *E. coli* DH5α treatments exhibited low frequencies of abnormal morphology, averaging approximately 8.4% ± 0.7%, 6.6% ± 1.15%, and 6.4% ± 0.3%, respectively, for bulbous hyphae. Together, these results indicate that metabolites produced by DC isolates strongly alter hyphal growth patterns and morphogenesis in *M. oryzae*.

### DC strains suppress *M. oryzae* growth through volatile-mediated mechanisms

Once *B. pumilus* DC strains were confirmed to exhibit direct antagonism against *M. oryzae* in dual-culture assays, we next evaluated whether these strains also produced volatile organic compounds (VOCs) that could inhibit fungal growth. VOC activity was assessed using a sealed stacking plate assay (Figure 5A), in which the bacteria and fungus shared the same headspace but did not physically contact each other [18]. Diameter growth was measured at 5 dpi to minimize the risk of cross-contamination between plates, including contamination from mature fungal spores or bacterial droplets. All three *B. pumilus* strains generated VOCs that reduced *M. oryzae* radial growth by approximately 30-45% compared with the untreated control (Figure 5B, C). Under the same conditions, the negative control *E. coli* DH5α did not measurably affect fungal growth. These results indicate that DC01, DC09, and DC13 release biologically active VOCs that suppress *M. oryzae* colony expansion. When combined with the dual-culture assays, these data demonstrate that the three *B. pumilus* isolates inhibit *M. oryzae* via both contact-dependent and volatile-mediated mechanisms, highlighting their potential as multifaceted biocontrol agents.

**Figure 5.**
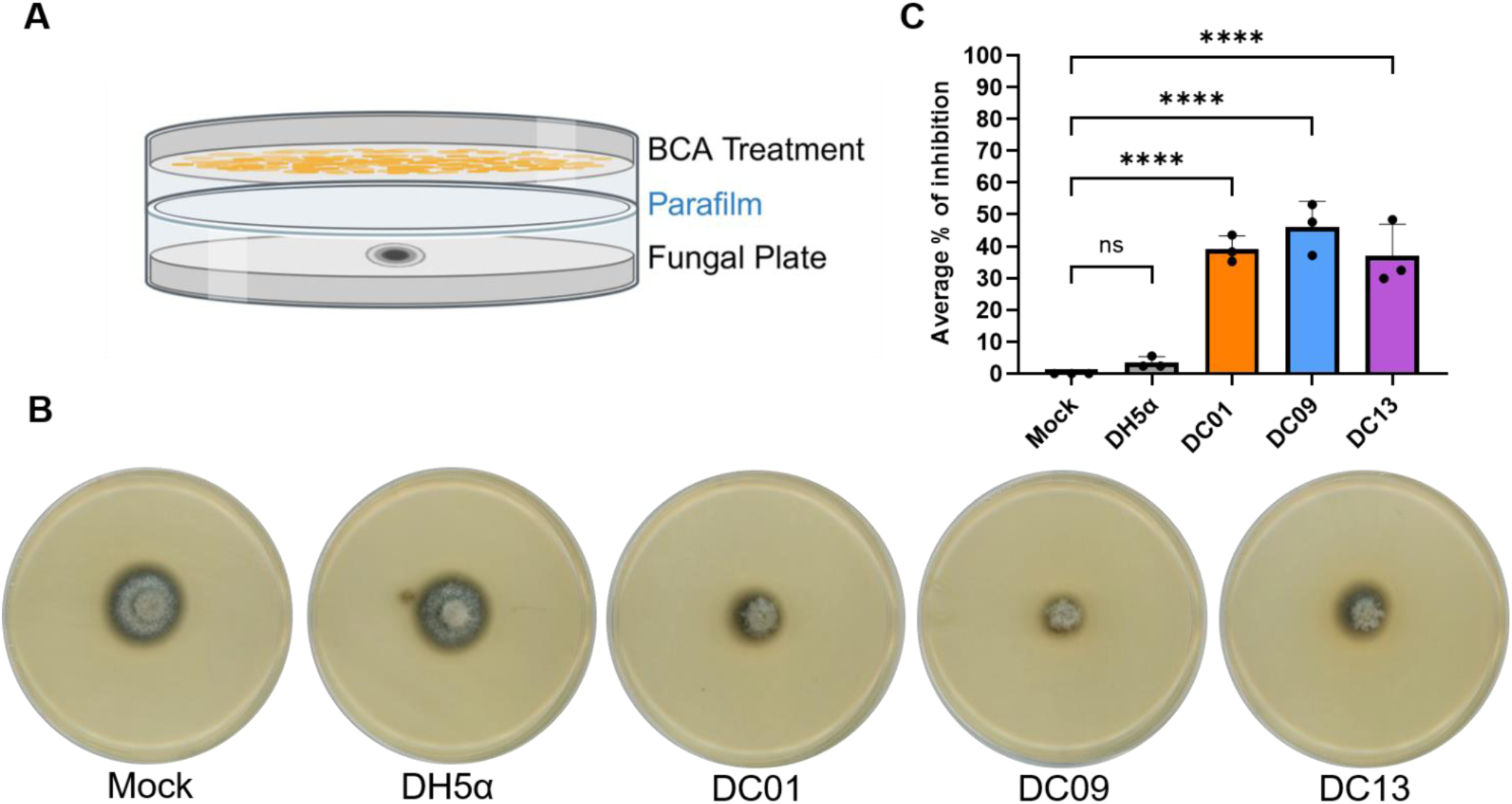
Volatility assay of DC strains against *M. oryzae. (*a) Diagram illustrating the volatile-mediated assay using a stacked plate method. (b) Fungal plates from the volatile-mediated assay after 5 dpi. (**c**) Quantification of the percentage of fungal growth inhibition in the presence of DCs and control samples (mock-water inoculation and *E. coli* DH5α). Bars represent the mean ± SD from three independent biological replicates (*n* = 3). Statistical significance was determined by one-way ANOVA followed by Dunnett’s multiple-comparison test against mock control. ns, not significant; \*\*\*\**p* < 0.0001.

### DC strains produce IAA and exhibit nitrogen fixation potential

To evaluate the PGP traits of the *B. pumilus* DC strains, we conducted a series of assays targeting key plant-beneficial functions. IAA production by *B. pumilus* DC strains was quantified using the Salkowski reagent method. The development of pink/red coloration when CFS is incubated with Salkowski’s reagent indicates the presence of indole compounds, proportional to IAA concentration. All three strains produced a clear pink-to-red signal and measurable IAA levels (Figure 6A). DC01, DC09, and DC13 synthesized 54.9 ± 0.52, 64.0 ± 0.15, and 46.3 ± 0.14 µg/mL IAA, respectively, while the positive-control strain *Azospirillum brasilense* Sp7 (JM01) reached 106.5 ± 0.12 µg/mL (Figure 6B). No color change or IAA signal was detected in the reagent-only blank.

**Figure 6.**
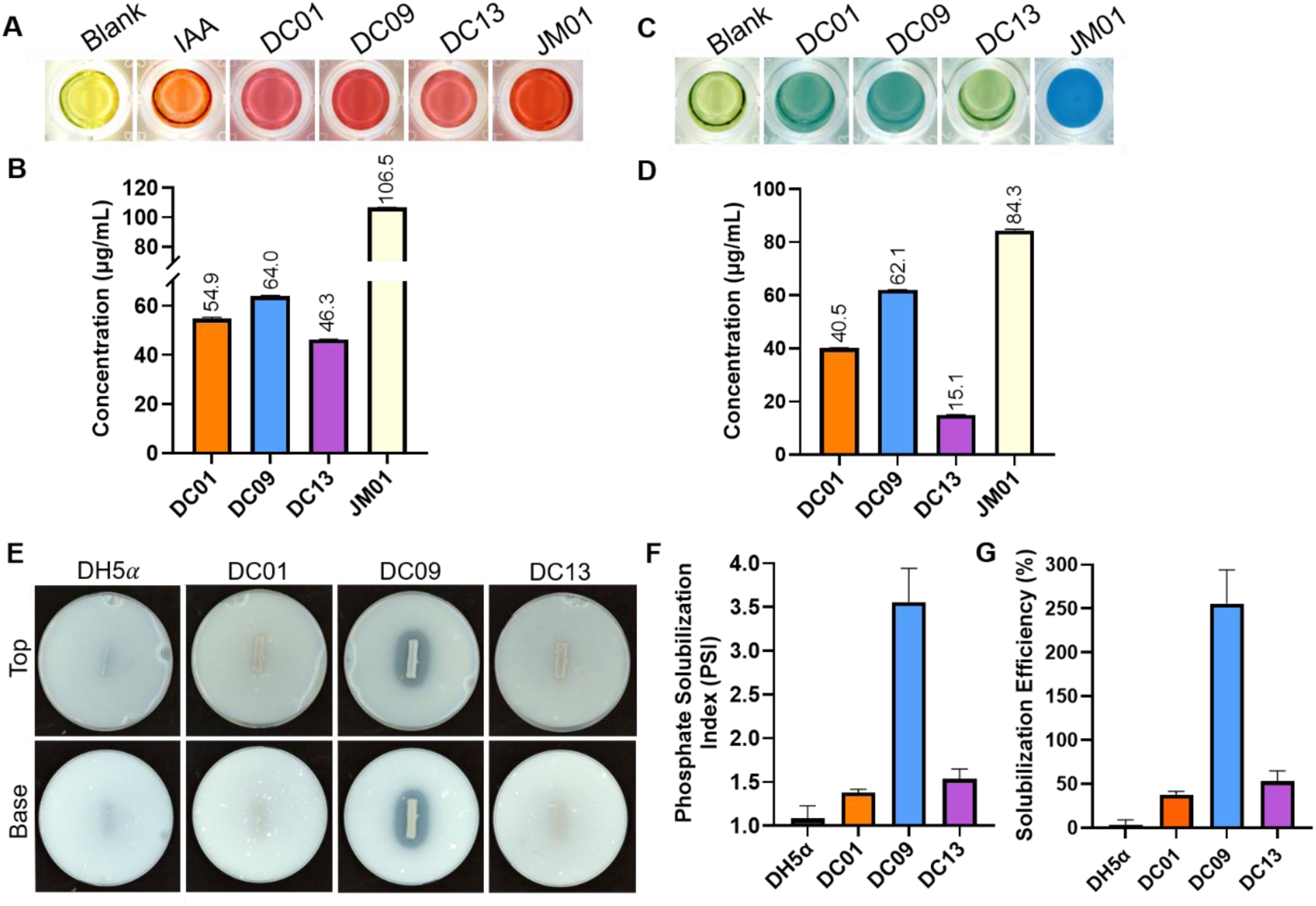
Plant growth-promoting traits of *B. pumilus* DC isolates. (**a-b**) IAA production was detected using Salkowski’s reagent; pink/red coloration indicates IAA synthesis. Values are expressed as µg/mL IAA equivalents. The 50 µg/mL IAA standard and the positive-control strain *A. brasilense* Sp7 (JM01) were included as positive controls, and the blank sample consisted of reagent only. (**c-d**) Nitrogen fixation potential is expressed as µg/mL of nitrogen-containing compounds produced in a nitrogen-free medium. (**e**) Qualitative phosphate solubilization on Pikovskaya’s medium plates, showing clear halos around bacterial colonies. (**f**) Phosphate Solubilization Index (PSI) for each strain. (**g**) Solubilization Efficiency (SE, %) of each strain. Bars represent mean ± SD from *n* = 3 biological replicates

Next, to evaluate the nitrogen-fixing potential of the DC strains, each isolate was cultured in nitrogen-free bromothymol blue (NFb) medium. DC01 and DC09 produced a more intense blue coloration than DC13, suggesting stronger metabolic activity under nitrogen-limited conditions (Figure 6C). Nitrogen-containing metabolite accumulation was subsequently quantified using a colorimetric OD610 assay. DC01, DC09, and DC13 produced 40.5 ± 0.07, 62.1 ± 0.03, and 15.1 ± 0.06 µg/mL fixed-nitrogen equivalents, respectively, compared with 84.3 ± 0.50 µg/mL for the positive control *A. brasilense* Sp7 (JM01) (Figure 6D). These results indicate that DC strains exhibit measurable nitrogen-fixing potential in NFb medium, together with IAA production, supporting their classification as PGPB.

### Phosphate solubilization differs among the *B. pumilus* isolates

To detect phosphate-solubilizing activity, *B. pumilus* DC strains were inoculated as lines on Pikovskaya’s agar, and halo formation was monitored along the bacterial streak. DC09 produced a pronounced clearing zone, whereas DC13 and DC01 generated smaller and less distinct halos. In contrast, the negative-control strain *E. coli* DH5α showed no visible halo (Figure 6A). Quantitative measurements revealed that DC09 exhibited the strongest phosphate-solubilizing activity, with a Phosphate Solubilization Index (PSI) of 3.55 ± 0.4 and a Solubilization Efficiency (SE) of 255% ± 38%, indicating robust mobilization of insoluble phosphate (Figure 6F, G). DC01 and DC013 exhibited low solubilization, with PSI values of 1.37 ± 0.04 and 1.53 ± 0.11, respectively, and SE values of 37.7% ± 3.8% and 53.33% ± 11%. By comparison, *E. coli* DH5α displayed only minimal solubilization (PSI = 1.08; SE = 8.33%), close to the basal level observed for the water control. Together, these results demonstrate that *B. pumilus* DC09 can solubilize inorganic phosphate and exhibits the highest phosphate-solubilizing potential among the tested strains.

### PEG-induced osmotic stress reveals enhanced tolerance in *B. pumilus* DC strains

To assess whether the *B. pumilus* DC strains possess traits associated with drought resilience, we evaluated their growth under osmotic stress induced by polyethylene glycol (PEG). Growth kinetics monitored over 20 h revealed that all three DC strains grew substantially faster in greater PEG concentrations than *E. coli* DH5α (Figure 7A-D). While growth progressively declined as PEG concentration increased, DC01, DC09, and DC13 continued to proliferate under both 20% and 30% PEG conditions, whereas *E. coli* DH5α exhibited severe growth inhibition. To further quantify osmotic stress tolerance, the area under the curve (AUC) was calculated and normalized to untreated controls to determine relative growth (%) (Figure 7E). At 10% PEG, DC01 showed a relative growth of 92.3% ± 2.34%, DC09 85.7% ± 2.23%, and DC13 90.2% ± 6.24%, whereas *E. coli* DH5α reached only 67.8% ± 6.28%. At 20% PEG, growth decreased across strains; however, the DC isolates maintained significantly greater tolerance, with DC01 at 56.4% ± 1.57%, DC09 at 53.1% ± 0.29%, and DC13 at 53.5% ± 3.41%, compared to *E. coli* DH5α at 15.3% ± 1.74%. Under 30% PEG, DC01 (30.1% ± 1.95%), DC09 (29.1% ± 2.99%), and DC13 (31.2% ± 1.3%) retained markedly higher growth, whereas *E. coli* DH5α dropped to 11.78% ± 0.10%. While slight variations were observed among DC strains, their overall responses were highly similar across conditions, indicating no clear differentiation in drought tolerance among the isolates. Together, these findings demonstrate that all *B. pumilus* DC strains exhibit strong and consistent tolerance to PEG-induced osmotic stress, as supported by both growth kinetics and AUC-based quantification.

**Figure 7.**
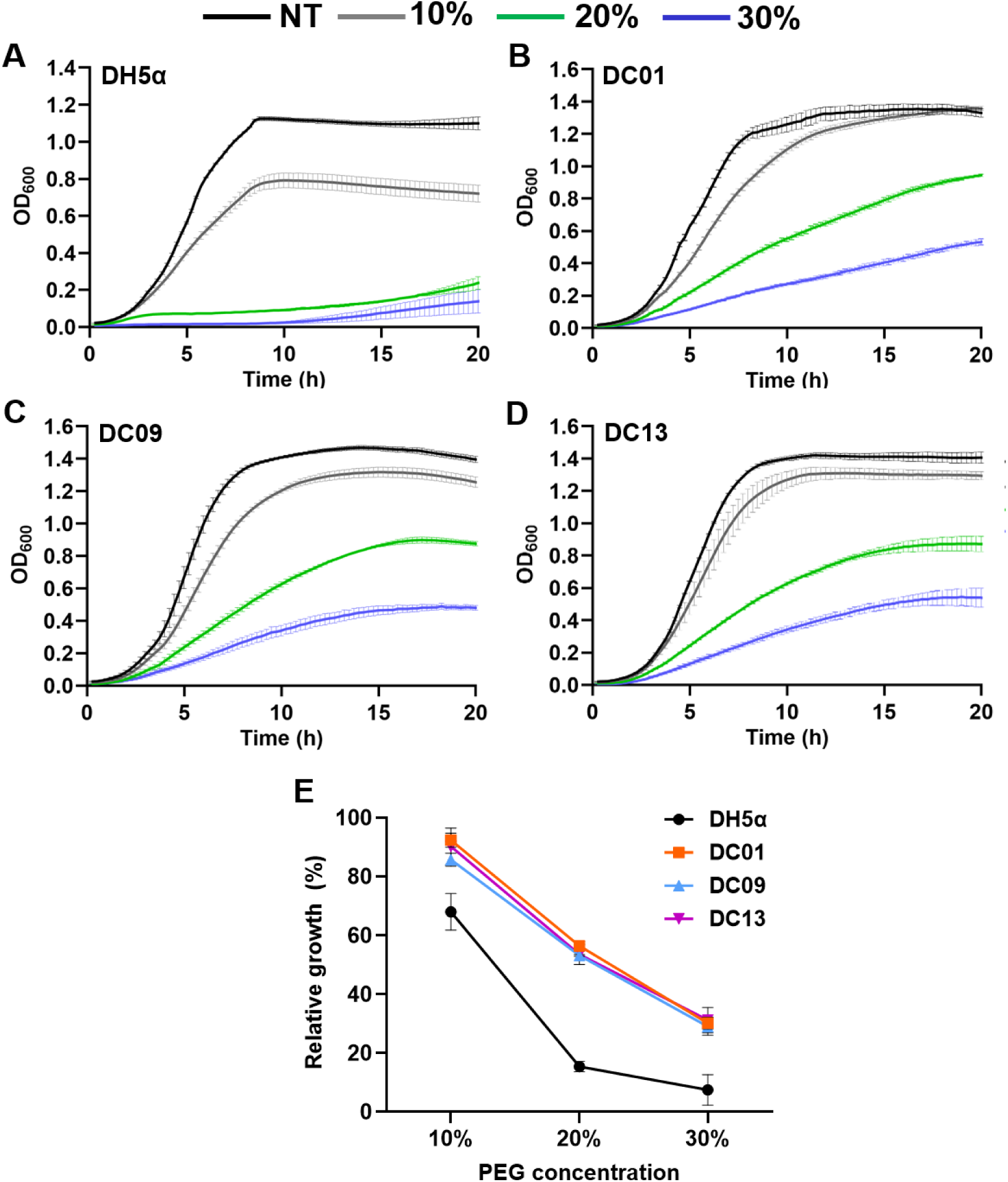
Drought tolerance of *B. pumilus* DC strains. (**a-d**) Growth curves of DC01, DC09, DC13, and *E. coli* DH5α cultured in LB medium supplemented with 0%, 10%, 20%, or 30% PEG 6000 to simulate osmotic stress. NT indicates the no-treatment control (0% PEG). Growth was monitored by OD₆₀₀ measurements over 20 h. (**e**) Relative bacterial growth (%) calculated from the area under the growth curve (AUC) and normalized to the corresponding NT condition for each strain. Values represent mean ± SD of three biological replicates. Statistical significance was determined by two-way ANOVA followed by Dunnett’s multiple-comparison test comparing each *B. pumilus* strain with *E. coli* DH5α at the same PEG concentration (*p* < 0.05).

### Soil inoculation with *B. pumilus* DC strains reduces rice blast severity and induces resistance

To evaluate whether the *B. pumilus* DC strains could reduce rice blast disease severity, we conducted a root-inoculation bioassay followed by foliar challenge with *M. oryzae* Guy11. Rice seedlings were root-treated with bacterial suspensions of DC01, DC09, or DC13 for 24 h and subsequently spray-inoculated with *M. oryzae* spores. Plants inoculated with the DC strains developed substantially fewer lesions than the mock and *E. coli* DH5α treatments (Figure 8A). While mock- and *E. coli* DH5α-treated leaves displayed extensive necrotic lesions characteristic of severe blast infection, leaves from DC-treated plants exhibited markedly reduced lesion density and overall diseased leaf area (Figure 8A). Quantification of disease severity further confirmed that all three DC strains significantly reduced blast symptoms compared with the control treatments (Figure 8B). These results demonstrate that a 24 h pre-treatment with the DC isolates induces a protective response that restricts pathogen development in rice leaves.

**Figure 8.**
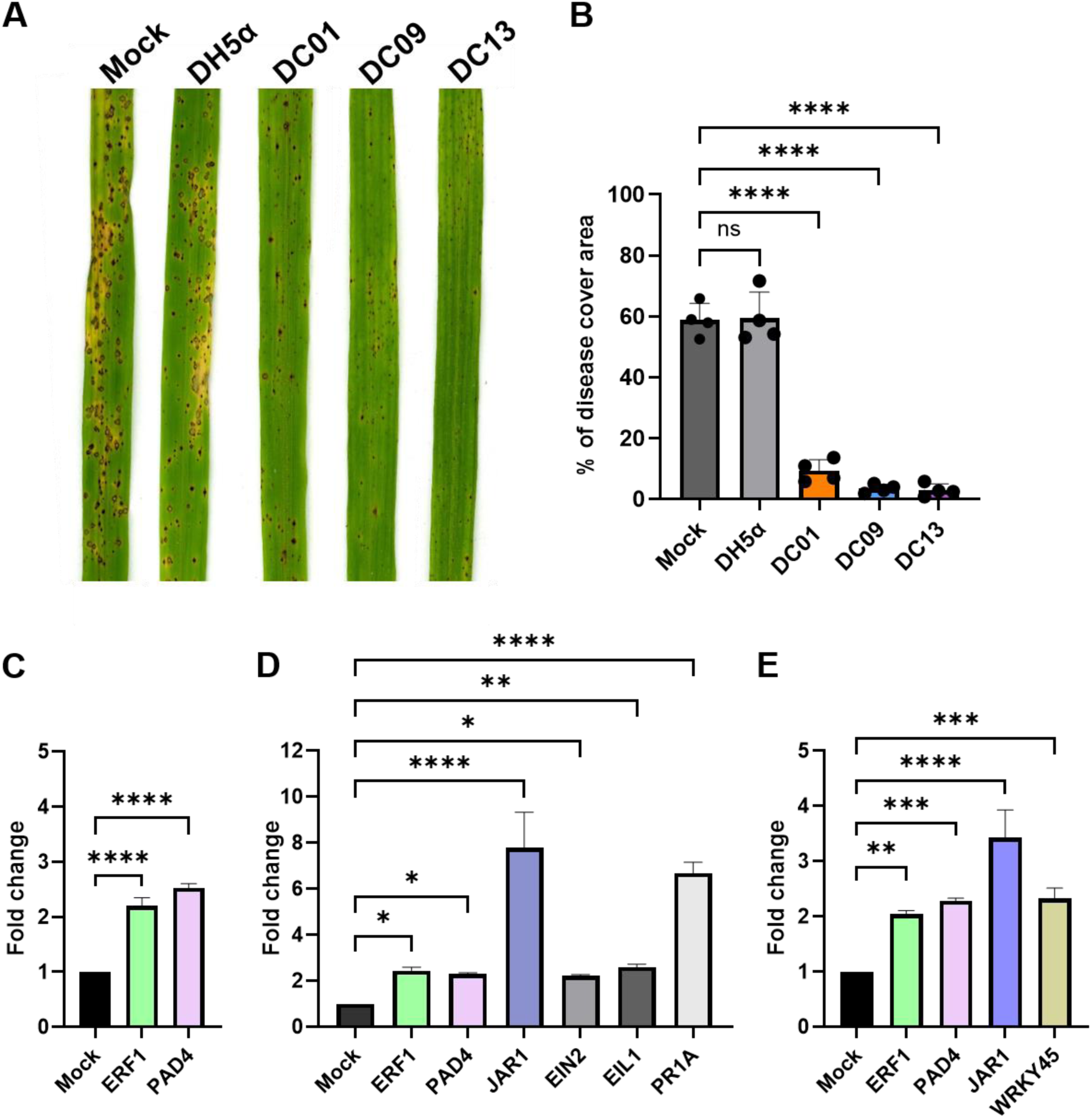
DC bacterial isolates induce systemic resistance in rice against *M. oryzae*. (**a**) Representative disease symptoms on CO39 rice leaves from mock- and DC-treated plants. Roots were inoculated with bacterial suspensions, and 24 h later, plants were spray-inoculated with *M. oryzae* (1 × 10^5^ spores/mL). (**b**) Disease severity was quantified as the percentage of leaf area covered by blast lesions. Bars represent mean ± SD with individual biological replicates shown as dots (*n* = 4). (**c-e**) Relative expression of defense-related genes in rice measured by qRT-PCR, normalized to GAPDH, and calculated using the ΔΔCt method. (c) DC01 (d) DC09 (e) DC13. Data are mean ± SD (*n* = 3 biological replicates). Statistical significance was determined by one-way ANOVA followed by Dunnett’s multiple comparison test against the mock control. *p* < 0.05 (\**), p <* 0.01 (**)*, p <* 0.001 (***), *p* < 0.0001 (****).

### Rice defense pathways are activated following DC strain inoculation

To determine whether the DC strains trigger early immune signaling consistent with ISR, we quantified the expression of a panel of JA-, ET-, SA-, and defense-associated marker genes 24 h after root inoculation with *B. pumilus* DC01, DC09, or DC13. The genes examined included *OsJAR1*, *OsERF1*, *OsEIN2*, *OsEIL2* (JA/ET signaling markers), OsWRKY45 and *OsPR1a* (SA-responsive markers), and *OsPAD4* (a defense-associated marker). All three DC strains altered the expression of at least a subset of these markers relative to non-inoculated controls, but the magnitude and pattern of induction differed among isolates (Figure 8C-E). DC09 produced the strongest overall transcriptional response, with notable increases in JA/ET-associated genes (*OsJAR1, OsERF1, OsEIN2, OsEIL2*) (Figure 8D). *OsJAR1* expression was also elevated in DC13-treated plants, but not in DC01-treated plants, suggesting strain-specific activation of JA-related signaling. DC01 and DC13 elicited more moderate or gene-specific expression changes, suggesting that early immune activation varies across strains (Figure 8C, E). SA-responsive genes showed modest induction across treatments. *OsWRKY45* and *OsPR1a* increased primarily in DC09-treated plants, while *OsERF1* and *OsPAD4* were the only markers consistently upregulated by all three isolates (Figure 8C-E). The shared induction of *ERF1* and *PAD4* suggests that these regulators participate in a common early response to root colonization. Because these transcriptional changes occurred in the absence of pathogen challenge, they reflect early immune signaling rather than infection-driven defense.

### DC strains exhibit broad-spectrum antagonistic activity

To assess whether the inhibitory activity of the DC strains extends across multiple taxa, we evaluated their antagonistic effects against selected fungal plant pathogens and bacterial human-associated pathogens. In dual-culture assays, all three *B. pumilus* DC strains inhibited the growth of agriculturally important fungi, although the magnitude of inhibition varied among strains and pathogens. DC01, DC09, and DC13 reduced the radial growth of *Botrytis cinerea* ME-1 by approximately 70%, 60%, and 55%, respectively, and of *Colletotrichum orbiculare* RH-18 by 20%, 30%, and 17%, as evidenced by restricted fungal expansion toward the bacterial streak (Table 2, Figure S1). These results indicate that the DC strains possess antifungal activity against multiple plant pathogens.

**Table 2.** Broad-spectrum antimicrobial activity of *B. pumilus* strains against phytopathogenic fungi and human-associated bacteria.

| Strains | DC01 | DC09 | DC13 |
| --- | --- | --- | --- |
| *Colletotrichum orbiculare* RH-18 | 20% | 30% | 17% |
| *Botrytis cinerea* ME-1 | 70% | 60% | 55% |
| *Micrococcus luteus* SK58 | 4.0 ± 1.9 mm | 4.2 ± 0.99 mm | 8.5 ± 1.34 mm |
| *Staphylococcus aureus* ATCC12600 | no | no | 5.2 ± 0.7 mm |
| *Escherichia coli* MG1655 | no | no | no |
| *Klebsiella pneumoniae* KP35 | no | no | no |
| *Pseudomonas aeruginosa* PAO1 | no | no | no |

Table note: Percent inhibition (%) indicates reduction of fungal radial growth relative to the untreated control. Inhibition against bacterial pathogens is reported as the diameter of the inhibition zone in millimeters (mm). Values are presented as mean ± standard deviation. “No” indicates no detectable inhibition under the assay conditions tested.

To evaluate antibacterial activity, DC01, DC09, and DC13 were tested against a panel of human-associated bacteria using cross-streak and spot-inoculation assays (Table 2), including the gram-positive species *Micrococcus luteus* SK58 and *Staphylococcus aureus* ATCC 12600, and the gram-negative species *E. coli* MG1655, *Klebsiella pneumoniae* KP35, and *Pseudomonas aeruginosa* PAO1. All three strains inhibited *M. luteus* SK58, producing measurable zones of inhibition ranging from 4.0 ± 1.9 mm (DC01) to 4.2 ± 0.99 mm (DC09), and a larger halo of 8.5 ± 1.34 mm for DC13. In contrast, inhibition of *S. aureus* ATCC 12600 was observed only for DC13 (5.2 ± 0.7 mm), whereas no detectable inhibition was observed for DC01 or DC09. No zones of inhibition were detected for any DC strain against the gram-negative bacteria tested (*E. coli* MG1655, *K. pneumoniae* KP35, or *P. aeruginosa* PAO1). Together, these data indicate that the DC strains, particularly DC13, exhibit stronger antibacterial activity against gram-positive taxa, consistent with the presence of predicted bacilysin, lipopeptide, and siderophore biosynthetic gene clusters in their genomes.

## Discussion

Rice blast caused by *M. oryzae* remains one of the most destructive diseases affecting rice production worldwide, underscoring the need for sustainable alternatives to chemical fungicides and the limited durability of resistance genes. In this study, we integrated comparative genomics, antagonistic assays, metabolite-based inhibition, and host transcriptional profiling to characterize three *B. pumilus* isolates as multi-mechanistic biological control agents against *M. oryzae*. Collectively, our findings support a model in which these strains suppress rice blast through both direct antifungal activity and host immune modulation, while also revealing substantial strain-specific variation in biocontrol-associated traits. Similar multifunctional biocontrol strategies have been widely associated with *B. pumilus* and related *Bacillus* spp., including antibiosis, volatile production, hydrolytic enzyme secretion, and ISR [20].

Whole-genome sequencing revealed that all three DC isolates harbor diverse NRPS and PKS BGCs, including clusters predicted to encode lipopeptides and siderophores commonly associated with *Bacillus*-mediated biocontrol. These genomic features are consistent with the experimentally observed antifungal and PGP traits and provide a genetic basis for the strain-specific differences observed among DC01, DC09, and DC13. Previous studies have shown that *B. pumilus* strains frequently encode biosynthetic pathways involved in the production of surfactins, fengycins, bacilysin, and other antimicrobial metabolites that contribute to fungal suppression and plant colonization [20]. The repeated detection of siderophore-associated clusters across all three genomes further supports their potential role in nutrient competition and rhizosphere fitness. Because the DC13 genome remains partially assembled, its complete biosynthetic repertoire may not yet be fully represented.

The DC isolates inhibited *M. oryzae* through multiple antagonistic mechanisms, including direct interaction, diffusible metabolites, and VOC-mediated suppression. In dual-culture assays, all three strains significantly reduced fungal radial growth, demonstrating strong antagonistic activity during direct interaction. However, clear strain-specific differences emerged in the cell-free supernatant assays. DC09 and DC13 retained strong inhibitory activity in the absence of living cells, whereas DC01 displayed minimal supernatant-mediated inhibition despite suppressing fungal growth in dual culture. The discrepancy between the dual-culture and CFS assays suggests that DC01-mediated inhibition may depend on direct bacterial-fungal interaction or localized metabolite production rather than freely diffusible compounds accumulated during growth in liquid culture. Similar observations have been reported in other *Bacillus* biocontrol systems, where strains displaying distinct lipopeptide signatures nevertheless exhibited comparable antifungal activity, suggesting that additional metabolites or interaction-dependent mechanisms may contribute substantially to pathogen suppression [28]. DC01 may therefore produce antifungal factors only in response to fungal proximity, or its inhibitory activity may be associated with cell-bound or surface-associated metabolites that are not efficiently recovered in the filtered supernatant. This distinction highlights mechanistic differences among the DC isolates despite their shared species classification.

An important observation was that antifungal activity persisted after heat treatment of the supernatants. Retention of inhibitory activity following heat inactivation suggests that at least part of the antifungal activity is mediated by heat-stable small molecules, consistent with *Bacillus*-derived cyclic lipopeptides and related secondary metabolites reported in previous studies [20, 29]. Cyclic lipopeptides such as fengycins, surfactins, and iturins are well known for their stability and membrane-disrupting activity against fungal pathogens [28–32]. Although the specific metabolites responsible for inhibition remain to be identified, the combination of predicted biosynthetic clusters and robust supernatant activity strongly supports a role for secreted secondary metabolites in the antagonistic activity of DC09 and DC13.

Microscopic analysis further demonstrated that DC supernatants substantially altered the morphology of *M. oryzae* hyphae. Exposure to supernatants from DC01, DC09, and DC13 induced bulbous swelling, altered hyphal polarity, apical deformation, and increased branching, with DC09 and DC13 producing the most pronounced structural abnormalities. Similar morphological alterations have previously been associated with antifungal lipopeptides produced by *Bacillus* spp., particularly members of the iturin and fengycin families, which disrupt fungal membrane integrity by permeabilizing membranes and destabilizing lipid bilayers [30–32]. These compounds have been reported to inhibit several phytopathogenic fungi, including *Botrytis cinerea*, *Fusarium* spp., and other filamentous fungi. In contrast, the comparatively weaker phenotype induced by DC01 is consistent with its limited inhibitory activity in the cell-free supernatant assays.

VOCs produced by DC isolates also contributed to fungal suppression. Physically separated VOC assays demonstrated that fungal growth inhibition could occur in the absence of direct contact or shared medium, indicating that volatile metabolites participate in antagonism. Similar VOC-mediated inhibition has been reported for several *Bacillus* species, including *B. pumilus*, in which volatile compounds contribute to the suppression of fungal phytopathogens in soil and rhizosphere environments [20, 33, 34]. The ability to inhibit pathogens through VOCs may be particularly advantageous under natural conditions, where volatile compounds can diffuse through soil pore spaces and influence microbial interactions at a distance. Although attempts were made to identify the volatile compounds produced by the DC isolates, these efforts were unsuccessful under the conditions tested. Future studies aimed at characterizing the volatile metabolite profiles of these strains will help clarify the contribution of VOCs to DC-mediated fungal inhibition.

Beyond *M. oryzae*, the DC isolates also displayed antagonistic activity against additional fungal and bacterial taxa, indicating that their inhibitory capacity is not restricted to a single pathogen. All three strains suppressed the growth of the phytopathogenic fungi *B. cinerea* and *C. orbiculare*, with varying degrees of inhibition observed among isolates. The DC isolates additionally inhibited the gram-positive bacteria *M. luteus*, while DC13 uniquely suppressed *S. aureus*. In contrast, no detectable inhibition was observed against the gram-negative bacteria tested. This differential activity spectrum is consistent with previous reports showing that *Bacillus*-derived antimicrobial compounds often display stronger activity against gram-positive organisms due to differences in cell envelope structure and membrane susceptibility [20]. Collectively, these findings suggest that DC isolates produce a diverse repertoire of antagonistic compounds with varying target specificity.

In addition to antifungal activity, all DC isolates displayed PGP traits, although these traits varied substantially among strains. Members of the genus *Bacillus* are widely recognized for their ability to enhance plant growth through phytohormone production, nutrient mobilization, and stress tolerance enhancement [13, 35]. In the present study, all three isolates produced IAA and tolerated osmotic stress, while DC09 uniquely displayed phosphate-solubilizing activity. These findings reinforce the concept that plant-beneficial traits are highly strain-dependent, even among closely related isolates. Previous studies similarly demonstrate that *B. pumilus* strains vary considerably in their expression of phosphate solubilization, phytohormone production, siderophore secretion, and stress adaptation traits [20].

Importantly, root inoculation with all three DC strains significantly reduced rice blast disease severity *in planta*, demonstrating that these isolates retain biocontrol activity under host-associated conditions. To investigate whether host immune activation contributed to disease suppression, we analyzed the expression of defense-associated genes involved in JA/ET signaling and SA-associated responses. All three DC strains altered host transcriptional responses relative to mock-treated plants, although the magnitude and composition of induction differed among isolates. A central observation was the consistent upregulation of *OsERF1* and *OsPAD4* across all DC treatments. In rice, *OsERF1* is associated with JA/ET-responsive defense signaling, whereas *OsPAD4* has been implicated in rice host–pathogen interactions and basal defense responses [36–38]. This suggests that root colonization by the DC isolates establishes an enhanced defensive state prior to pathogen challenge. Notably, DC09 induced the strongest activation of JA/ET-associated genes, including *OsERF1*, *OsJAR1*, *OsEIN2*, and *OsEIL2*. This transcriptional profile is consistent with ISR-like activation and aligns with previous studies demonstrating that *Bacillus* spp. can stimulate JA/ET-mediated defense pathways in plants [20, 24–26]. In contrast, DC01 and DC13 induced more moderate transcriptional responses, suggesting that although all three strains engage host defense pathways, the magnitude and composition of immune activation differ between isolates.

Taken together, our genomic, *in vitro*, and *in planta* analyses support a model in which the DC isolates suppress rice blast through multiple complementary mechanisms, including diffusible antifungal metabolites, VOC-mediated inhibition, disruption of fungal development, and activation of host defense pathways. Among the isolates, DC09 and DC13 emerged as the most functionally integrated strains, combining strong metabolite-mediated antagonism with broad host transcriptional activation. More broadly, the substantial phenotypic differences observed among these closely related isolates highlight the importance of evaluating *Bacillus* strains individually rather than relying solely on species-level classifications when developing BCAs. To our knowledge, this study represents one of the most comprehensive characterizations of strain-specific antagonistic and plant-associated traits in *B. pumilus* isolates active against *M. oryzae*, integrating comparative genomics, metabolite-associated inhibition, fungal morphogenesis, VOC-mediated antagonism, and host defense activation.

Future work should focus on identifying the metabolites responsible for antifungal activity, linking specific BGCs to inhibitory phenotypes, further characterizing the molecular basis of host immune activation, and evaluating the efficacy of the DC isolates under field conditions. Nonetheless, the present study demonstrates that the DC isolates, particularly DC09 and DC13, represent promising candidates for sustainable management of rice blast disease and further illustrate the potential of *B. pumilus* as a versatile platform for agricultural biocontrol.

## Materials and methods

### Microbial strains and culture conditions

The *B. pumilus* DC strains used in this study were recovered from various topsoil samples collected in the Gainesville, Florida area (29.643280° N, -82.331693° W; 29.624527° N, -82.361428° W; and 29.6241345° N, -82.3638663° W). Bacterial strains were routinely cultured and maintained in Luria-Bertani (LB) broth containing 10 g/L tryptone, 5 g/L yeast extract, and 5 g/L NaCl. Medium was autoclaved at 121 °C and 15 psi for 20 min prior to use. For routine culturing, bacterial strains were grown overnight in 3 mL LB broth at 30 °C with shaking at 200 revolutions per minute (rpm). Long-term stocks were maintained in 20% glycerol at −80 °C.

The fungal strain *Magnaporthe oryzae* Guy11 was used throughout this study and maintained on complete medium (CM). CM consisted of 50 mL of 20× nitrate salts, 1 mL trace elements, 10 g D-glucose, 2 g peptone, 1 g yeast extract, 1 g casamino acids, and 1 mL vitamin solution per liter of distilled water. The pH was adjusted to 6.5 using 5 M NaOH. For a solid medium, 15 g/L agar was added prior to autoclaving. Long-term fungal stocks were stored on sterile Whatman filter paper at −20 °C.

Additional fungal strains used in antagonism assays, including *Botrytis cinerea* RH-18 and *Colletotrichum orbiculare* ME-1, were kindly provided by Dr. Jeffrey Rollins’ laboratory. Additional bacterial strains used in antagonism assays, including *Micrococcus luteus* SK58, *Staphylococcus aureus* ATCC 12600, *Escherichia coli* MG1655, *Klebsiella pneumoniae* KP35, and *Pseudomonas aeruginosa* PAO1 were cultured in LB broth at 37 °C with shaking at 200 rpm.

### Plant growth conditions

Seeds of *Oryza sativa* cultivar CO39 were dehusked and surface sterilized by immersion in 70% ethanol for 1 minute, rinsed with sterile water for 1 minute, followed by incubation in 5% bleach for 1 h. Seeds were rinsed three times with sterile water, dried on sterile paper towels, and planted in Magenta boxes containing 100 mL of half-strength Murashige and Skoog (½ MS) medium (2.25 g MS salts, 30 g sucrose, and 0.5 g MES per liter of distilled water). The pH was adjusted to 5.8 using KOH, followed by the addition of 3 g/L gelrite. The medium was autoclaved at 121 °C for 25 minutes, and 100 mL was dispensed aseptically into each autoclaved Magenta box. Plants were grown for 10 days at 25 °C under a 12-h light/dark photoperiod in a growth chamber. Ten-day-old seedlings were transplanted into 4-inch surface-sterilized pots containing autoclaved soil (Pro Mix HP Mycorrhizae: Michigan Peat Baccto General Purpose Potting Soil) and maintained in the greenhouse.

### Whole-genome sequencing, assembly, and annotation of bacterial isolates

Bacterial isolates were grown overnight in 5 mL of fresh LB medium at 30 °C while shaking at 200 rpm. Genomic DNA was extracted from bacterial isolates using the AllPrep DNA/RNA/Protein Kit (Qiagen) according to the manufacturer’s protocol. DNA concentration and purity were assessed spectrophotometrically, and the samples were submitted to SeqCenter (Pittsburgh, PA) for 16S rRNA gene amplicon sequencing (V1 and V4 primers) and whole-genome sequencing (WGS) using the Illumina platform. Taxonomic identification was performed using 16S rRNA gene sequences and BLASTn against the NCBI nucleotide database. For WGS, raw Illumina reads were quality-checked using FastQC [39] and then processed with the BBTools suite [40] to remove adapters and phiX contamination. Cleaned reads were assembled de novo using SPAdes v3.15.5 [41], and assemblies were polished with two iterations of Pilon [42]. Draft genome assemblies were uploaded to the Bacterial and Viral Bioinformatics Resource Center (BV-BRC) [43] for species-level identification and functional annotation using RASTtk. Biosynthetic gene clusters were predicted using antiSMASH with relaxed detection strictness and all additional features enabled, and predicted coding sequences within secondary metabolite regions were assigned putative functions by BLASTp analysis [27]. Phylogenetic relationships were inferred using a codon-based tree constructed in BV-BRC with RAxML based on 100 single-copy genes, including the isolates from this study and 49 reference *Bacillus pumilus* genomes; trees were visualized and annotated using the interactive Tree of Life (iTOL) web interface [44].

### Dual culture assay

To assess antagonistic activity, bacterial overnight cultures were adjusted to an OD_600_ of 0.5 using sterile water. A UV-sterilized 5 mm core borer, forceps, and rod were used to collect agar plugs from the actively growing edge of 7-day-old *M. oryzae* cultures. Fungal plugs were placed approximately 1.5 cm from the center of a CM agar plate, ensuring mycelial contact with the media. Opposite each fungal plug and equidistant from the center, 5 µL of normalized bacterial culture was spotted onto the plate. As a negative control, sterile water was applied in place of the bacterial inoculum. Plates were incubated at 25 °C in the dark. Unless otherwise stated, fungal growth was assessed at 10 dpi. Fungal colony diameters and radii were measured and averaged for each replicate. The percentage of growth inhibition was calculated using the formula: Inhibition (%) = [(average control - average experimental)/average control] × 100. This applies to the radius and diameter of the fungal colony.

### Cell-free culture supernatant (CFS) inhibition assay

To evaluate the effect of bacterial metabolites on *M. oryzae* growth, we prepared cell-free supernatants (CFS) from the *B. pumilus* DC strains. Each DC isolate was grown in tryptic soy broth (TSB; Difco) at 30 °C with shaking at 180 rpm for 24 h. Cultures were then centrifuged to remove bacterial cells, and the resulting supernatants were carefully collected and filter-sterilized through 0.22 µm syringe filter (VWR) to obtain CFS. TSB medium was used as a mock control. For heat-inactivation assays, CFS were incubated at 95 °C for 10 min before being incorporated into the medium. For the plate assay, CFSs or sterile TSB (control) were mixed with CM at a 1:6 (v/v) ratio, thoroughly mixed, and poured into Petri dishes. After solidification, a 5 mm mycelial plug of *M. oryzae* strain Guy11 was placed in the center of each plate. Plates were incubated at 25 °C in the dark for 5 days. At the end of the incubation period, colony diameters were measured, and plates were scanned for documentation. The percentage of growth inhibition relative to the mock treatment was calculated using the same formula applied for the dual-culture assay.

### Microscopic analysis of fungal development

To further assess the effect of bacterial metabolites on fungal hyphal morphology, microscopic analysis was performed on CM agar and CM supplemented with CFS or TSB (control) at a 1:6 (v/v) ratio. Briefly, 250 µL of medium was applied to autoclaved microscope slides and allowed to dry prior to inoculation. A 5 µL spore suspension (5 × 105 spores/mL) of *M. oryzae* was applied to each end of the agar smear. Slides were incubated at 25 °C in humid chambers under a 12 h light/12 h dark photoperiod and imaged at 5 dpi using a Zeiss Axio Observer 7 microscope. For Calcofluor White staining, fungal growth was treated with one drop of Calcofluor White (Sigma) and one drop of 10% KOH, incubated for 1 min, and imaged immediately. Images were processed using ZEN Microscopy software v3.13 (Zeiss).

### Volatile-mediated inhibition assay

Volatile inhibition of *M. oryzae* by bacterial strains was assessed using a sealed dual-plate system. Bacterial cultures were grown and normalized to an OD_600_ of 0.5 using the same procedure as described for the dual culture assay. CM agar plates were inoculated with 5 µL of normalized bacterial culture and spread evenly using sterile glass beads. For the negative control, sterile water was used in place of the bacterial culture. Plates were incubated overnight at 28 °C in the dark to allow bacterial growth. Fungal plugs were obtained from the growing edge of 7-day-old *M. oryzae* cultures using sterilized tools and aseptically transferred to the center of fresh CM plates. The pre-inoculated bacterial plate (lid removed) was inverted and placed on top of the fungal plate, forming a sealed volatile chamber using parafilm. The assembled plates were incubated for 5 days at 25 °C in the dark. Following incubation, fungal growth was imaged, and colony diameter was measured in two perpendicular directions and averaged. The percentage of inhibition was calculated using the formula described in the dual-culture assay.

### Indole-3-Acetic Acid (IAA) production quantification

To quantify IAA production, 3 mL cultures of half-strength TSB supplemented with 0.5% (w/v) L-tryptophan were inoculated with bacterial strains normalized to an OD_600_ of 0.5. Cultures were incubated with shaking at 200 rpm for two days, beginning at 37 °C for the first 24 h and then shifted to 28 °C for the remaining 48 h. IAA production was assessed at 72 h. At this time, 500 µL of culture was centrifuged at 10,000 rpm for 10 minutes, and 50 µL of the resulting supernatant was transferred to a 96-well plate. Control conditions included a negative control containing 50 µL of 0.5% L-tryptophan-supplemented 1/5 TSB, a chemical positive control made by mixing 25 µL of 0.5% L-tryptophan TSB with 25 µL of 100 µg/mL IAA stock solution, and a biological positive control using *Azospirillum brasilense* (kindly provided by Dr. Jean-Michel Ané), a known IAA-producing strain. To each well, 100 µL of freshly prepared Salkowski reagent (50 mL of 35% perchloric acid mixed with 1 mL of 0.5 M FeCl_3_) was added, and samples were incubated in the dark at room temperature for 30 minutes [45, 46]. Color development was visually documented, with IAA-positive samples turning pink. Absorbance was measured at 536 nm using a microplate reader, and IAA concentrations were calculated using the standard curve trendline equation*: y = 0.0069x − 0.0013*, where *y* is the absorbance at 536 nm and *x* is the IAA concentration (µg/mL).

### Phosphate solubilization capacity

Bacterial cultures were prepared by inoculating 3 mL of LB broth with the bacterial strain of interest and incubating for 24 h at 30 °C with shaking at 200 rpm. Cultures were normalized to an OD600 of 0.5 using sterile distilled water, and cells were washed with water to remove any trace of LB. Phosphate solubilization was assessed using Pikovskaya’s agar (PVK; 10 g D-glucose, 5 g Ca_3_(PO_4_)_₂_, 0.5 g (NH_4_)_₂_SO_4_, 0.2 g KCl, 0.1 g MgSO_4_, trace amounts (∼0.001 g each) of MnSO_4_ and FeSO_4_, and 0.5 g yeast extract per liter) [46, 47]. A 5 µL aliquot of normalized culture was spotted at the center of PVK plates and streaked into a 3 cm line. Plates were incubated at 37 °C for seven days. After incubation, phosphate solubilization was evaluated by the formation of clear halos around bacterial growth. Plates were imaged, and halo and colony diameters were measured. Phosphate solubilization was quantified by calculating the Phosphate Solubilization Index (PSI) and Solubilization Efficiency (SE). PSI was calculated as: (Halo diameter + Colony diameter)/Colony diameter. SE was calculated as: SE (%) = [(Halo diameter - Colony diameter)/Colony diameter] x 100

### Nitrogen fixation assay

Nitrogen-free bromothymol blue (NFb) medium was prepared by combining the following components per liter of distilled water: 5 g malic acid, 0.5 g K_2_HPO_4_, 0.2 g MgSO_4_, 0.1 g NaCl, 0.02 g CaCl_2_·2H_2_O, 2 mL micronutrient solution (containing 0.04 g CuSO_4_·5H_2_O, 0.12 g ZnSO_4_·7H_2_O, 1.40 g H3BO3, 1.0 g Na_2_MoO_4_·2H_2_O, and 1.175 g MnSO_4_·H_2_O; brought to 1 L and stored at 4°C), 2 mL bromothymol blue solution (5 g bromothymol blue in 0.2 M KOH), 4 mL Fe-EDTA solution (16.4 g Fe-EDTA per liter), 1 mL vitamin solution (10 mg biotin and 20 mg pyridoxine in 100 mL distilled water), and 4.5 g KOH, the pH was set to 6.5 using KOH [48, 49]. After autoclaving, a 50 mL working stock was prepared with 49.833 mL of NFb medium and 166 µL of sodium lactate solution and stored at 4 °C.

To assess nitrogen fixation, 3 mL of overnight bacterial cultures were diluted in LB to an OD_600_ of 0.5. In a sterile 96-well plate, three columns were filled with 180 µL of NFb medium, and 20 µL of normalized bacterial culture was added to each well. *A. brasilense* was included as a biological positive control under the same conditions. Three wells containing 200 µL of uninoculated NFb medium served as negative controls. The plate was incubated at 37 °C for 24 h with shaking at 200 rpm. Absorbance was measured at 610 nm using a microplate reader. Color changes from green to blue were recorded as indicative of ammonium ion production. Fixed nitrogen concentration was determined using a standard curve and trendline equation, as reported in [49].

### Drought tolerance assay

Drought tolerance of bacterial isolates was assessed through culturing in liquid LB medium supplemented with 10%, 20%, and 30% (w/v) polyethylene glycol (PEG 6000) to simulate osmotic stress. Overnight bacterial cultures were grown in LB and normalized to an OD_600_ of 0.5. A 1:100 dilution of each culture was prepared in PEG-supplemented LB. In a sterile 96-well plate, 180 µL of each PEG-containing medium was aliquoted per well, and 20 µL of the diluted bacterial suspension was added. Control wells contained uninoculated LB medium. *E. coli* DH5α was included as a non-tolerant control. Plates were incubated at 37 °C with continuous shaking, and bacterial growth was monitored by measuring optical density at 600 nm (OD_600_) over a 20 h period using a microplate reader. Growth curves were generated over time, and total growth for each condition was quantified by calculating the area under the curve using Riemann sum approximation.

### Plant inoculations and infections

Overnight bacterial cultures were prepared and normalized to an OD600 of 0.1. Cultures were centrifuged at 10,000 × g for 3 minutes, washed twice with sterile water, and resuspended in 1 mL of sterile water. This suspension was diluted in 9 mL of sterile water to obtain a final inoculum. Ten-day-old rice seedlings were submerged in the bacterial suspension for 20 minutes, then transferred to sterile soil-filled 4-inch pots. Control plants were mock-inoculated using sterile water. At three weeks post-germination, plants were re-inoculated with 20 mL of the normalized bacterial culture applied directly to the soil. After 24 h, a *M. oryzae* spore suspension (1 × 10^5^ conidia/mL in 0.2% gelatin) was sprayed onto the leaves of each plant. Plants were covered with clear plastic bags to maintain high humidity and incubated at 25 °C under a 12-h light/dark cycle in a growth chamber. After two days, bags were partially opened to gradually reduce humidity. Five days post-inoculation, leaves were harvested, imaged, and disease severity was quantified as the percentage of diseased leaf area using ImageJ software [50].

### Gene expression analysis

Total RNA was extracted from rice leaf samples using the RNeasy Plant Mini Kit (Qiagen) according to the manufacturer’s protocol. RNA concentration and purity were measured using a spectrophotometer, and RNA integrity was confirmed via agarose gel electrophoresis. DNA contamination was removed using DNase I treatment. First-strand complementary DNA (cDNA) was synthesized from 1 µg of total RNA in a final volume of 20 µL using the QuantaBio reverse transcription kit with oligo-dT primers, following the manufacturer’s instructions.

Quantitative PCR was performed using QuantaBio PerfeCTa SYBR Green FastMix in a Bio-Rad CFX96 thermal cycler with gene-specific primers. Each reaction was carried out in a total volume of 10 µL, including 2 µL of diluted cDNA, 0.5 µM of each primer (Table S2), and 1× SYBR Green master mix. The thermal cycling conditions were as follows: initial denaturation at 95 °C for 2 minutes, followed by 40 cycles of 95 °C for 15 seconds and 60 °C for 30 seconds. A melt curve analysis was included to confirm the specificity of amplification. GAPDH was used as the reference gene for normalization. Relative gene expression levels were calculated using the ΔΔCt method, and expression was reported as fold change relative to uninoculated control plants. All reactions were performed in technical triplicate, and results were averaged across three biological replicates.

### Statistical analysis

Statistical analyses and graphing were performed using GraphPad Prism v10.6.1 (GraphPad Software, Boston, MA, USA). Unless otherwise indicated, experiments were conducted using three independent biological replicates, each containing three technical replicates. Statistical significance was determined using one-way or two-way ANOVA followed by Dunnett’s or Tukey’s post hoc multiple comparison tests, as appropriate (*p* < 0.05). Error bars represent the standard deviation (SD) of biological replicates.

## Supplementary Materials

Figure S1: Antagonistic activity of *Bacillus* DC strains against *B. cinerea* and *C. orbiculare.*; Table S1: List of PCR primers used for qRT-PCR.

## Author Contributions

Conceptualization, J.F.; methodology, J.F., D.M.C, L.E.K., T.R.J., G.L.E., N.P., and R.E.K.; writing, J.F., D.M.C., G.L.E., L.E.K., and R.E.K.; supervision, J.F, D.M.C. All authors have read and agreed to the published version of the manuscript.

## Institutional Review Board Statement

Not applicable.

## Informed Consent Statement

Not applicable.

## Data Availability Statement

Sequencing data generated in this study have been depossited in the NCBI Sequence Read Archive (SRA) under the BioProject accession number PRJNA1480024.

## Acknowledgments

We thank members of the Fernandez laboratory for their constructive feedback and helpful discussions. We also thank the undergraduate students in the laboratory who indirectly supported this work by assisting with lab organization and maintaining supplies. We are grateful to Daniel Czyz and members of the Czyz laboratory for helpful edits and valuable feedback on the manuscript.

## Abbreviations

The following abbreviations are used in this manuscript:

PGPB: plant growth-promoting bacteria
IAA: Indole-3-acetic acid
ISR: Induced systemic resistance
VOC: Volatile organic compounds
BCA: Biocontrol agent
BGC: Biosynthetic gene clusters
SA: Salicylic acid
JA: Jasmonic acid
ET: Ethylene
T3PKS: Type III polyketide synthases
NRPS: Non-ribosomal peptide synthetase
CFS: Cell-free supernatant
Days: post-inoculation
CM: Complete media
TSB: Tryptic soy broth
NFb: Nitrogen-free bromothymol blue
PVK: Pikovskaya medium
SE: Solubilization Efficiency
PSI: Phosphate solubilization index
PEG: Polyethylene glycol
MS: Murashige and Skoog
LB: Luria-Bertani

