## Supplemental Figure 1 and Table 1 for "Soil-derived *Bacillus pumilus* strains demonstrate antagonistic activity against *Magnaporthe oryzae* and multiple plant growth-promoting traits"

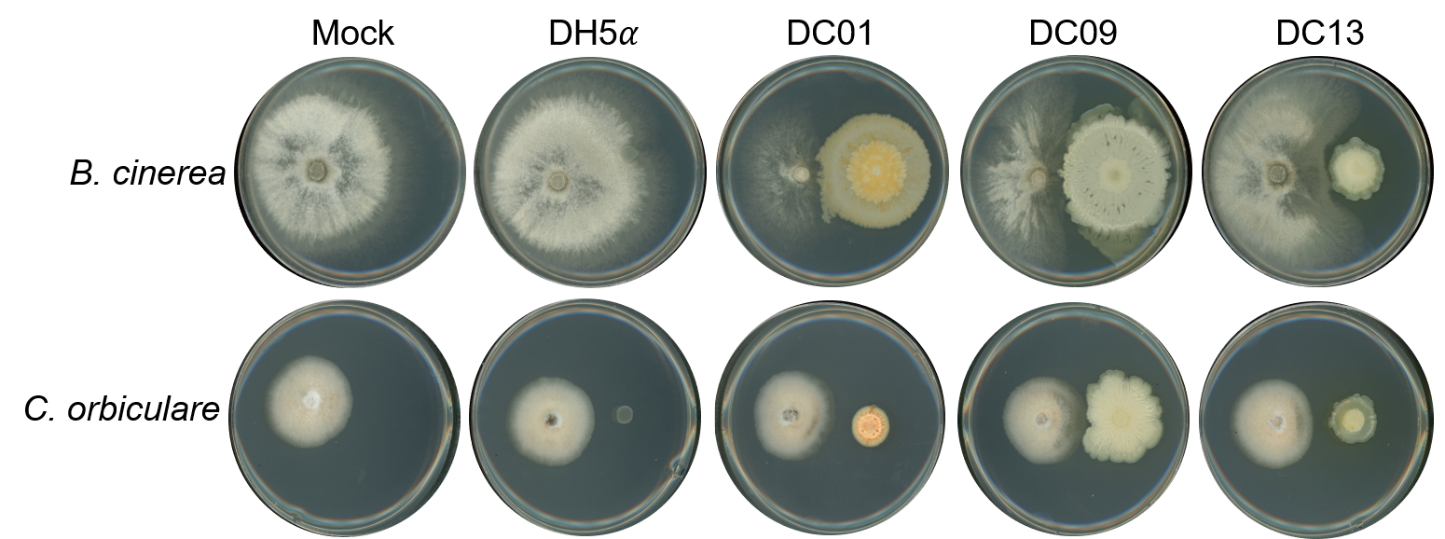

**Supplemental Figure 1**. Antagonistic activity of *Bacillus* DC strains against *B. cinerea* and *C. orbiculare.* Dual culture assay plates showing fungal growth inhibition by DC strains compared to mock and DH5$\alpha$ controls. Plates were incubated at 25 °C, and images were taken at 5 dpi.

**Supplemental Table 2**. List of PCR primers used for qRT-PCR.

| **Names** | **Primers Sequence (5’-3’)** | **Source** |
| --- | --- | --- |
| PAD4 | Forward: GCACAAGTTTGAGCGCCATT  Reverse: ACTCCTTCACCCACAGGGAGTA | [1] |
| WRKY45 | Forward: CGGGTAAAACGATCGAAAGA  Reverse: TTTCGAAAGCGGAAGAACAG | [1] |
| JAR1 | Forward: TCTCCCCAGCCTTAACCGTA  Reverse: CTAAACGCGACGACAAACCC | [2] |
| EIL1 | Forward: ACAATGCCACGATCATGGAG  Reverse: TCAGTAGTACCAATTCGAGC | [1] |
| PR1 | Forward: CGTCTTCATCACCTGCAACTACTC  Reverse: CATGCATAAACACGTAGCATAGC | [1] |
| ERF1 | Forward: CATATCACCTTGACGCCCCA  Reverse: ACCCTCACAAACTCACTCGG | [2] |
| EIN2 | Forward: CAAGGAACCAGTGACAACCA  Reverse: GCAGTCGTCTCCGCAGTTAG | [3] |
| GAPDH | Forward: AAGCCAGCATCCTATGATCAGATT  Reverse: CGTAACCCAGAATACCCTTGAGTTT | [4] |

1.	Ke Y, Liu H, Li X, Xiao J, Wang S. 2014. Rice OsPAD4 functions differently from *Arabidopsis* AtPAD4 in host-pathogen interactions. The Plant Journal 78:619-631. <https://doi.org/10.1111/tpj.12500>.

2.	Spence C, Alff E, Johnson C, Ramos C, Donofrio N, Sundaresan V, Bais H. 2014. Natural rice rhizospheric microbes suppress rice blast infections. BMC Plant Biol 14:130. <https://doi.org/10.1186/1471-2229-14-130>.

3.	Duan C, Yu J, Bai J, Zhu Z, Wang X. 2014. Induced defense responses in rice plants against small brown planthopper infestation. The Crop Journal 2:55-62. <https://doi.org/10.1016/j.cj.2013.12.001>.

4.	Chen X, Laborda P, Dong Y, Liu F. 2020. Evaluation of suitable reference genes for normalization of quantitative real-time PCR analysis in rice plants under *Xanthomonas oryzae* pv. *oryzae* infection and melatonin supplementation. Food Production, Processing and Nutrition 2:21. <https://doi.org/10.1186/s43014-020-00035-9>.
